# MurA escape mutations uncouple peptidoglycan biosynthesis from PrkA signaling

**DOI:** 10.1101/2021.09.09.459578

**Authors:** Sabrina Wamp, Patricia Rothe, Gudrun Holland, Sven Halbedel

## Abstract

Gram-positive bacteria are protected by a thick mesh of peptidoglycan (PG) completely engulfing their cells. This PG network is the main component of the bacterial cell wall, it provides rigidity and acts as foundation for the attachment of other surface molecules. Biosynthesis of PG consumes a high amount of cellular resources and therefore requires careful adjustments to environmental conditions.

An important switch in the control of PG biosynthesis of *Listeria monocytogenes*, a Gram-positive pathogen with a high infection fatality rate, is the serine/threonine protein kinase PrkA. A key substrate of this kinase is the small cytosolic protein ReoM. We have shown previously that ReoM phosphorylation regulates PG formation through control of MurA stability. MurA catalyzes the first step in PG biosynthesis and the current model suggests that phosphorylated ReoM prevents MurA degradation by the ClpCP protease. In contrast, conditions leading to ReoM dephosphorylation stimulate MurA degradation. How ReoM controls degradation of MurA and potential other substrates is not understood. Also, the individual contribution of the ∼20 other known PrkA targets to PG biosynthesis regulation is unknown.

We here present *murA* mutants which escape proteolytic degradation. The release of MurA from ClpCP-dependent proteolysis was able to constitutively activate PG biosynthesis and further enhances the intrinsic cephalosporin resistance of *L. monocytogenes*. This activation required the RodA3/PBP B3 transglycosylase/transpeptidase pair as additional effectors of the PrkA signaling route. One *murA* escape mutation not only fully rescued an otherwise non-viable *prkA* mutant during growth in batch culture and inside macrophages but also overcompensated cephalosporin hypersensitivity. Our data collectively indicate that the main purpose of PrkA-mediated signaling in *L. monocytogenes* is control of MurA stability during extra- and intracellular growth. These findings have important implications for the understanding of PG biosynthesis regulation and β-lactam resistance of *L. monocytogenes* and related Gram-positive bacteria.

**Author Summary:** Peptidoglycan (PG) is the main component of the bacterial cell wall and many of the PG synthesizing enzymes are antibiotic targets. We previously have discovered a new signaling route controlling PG production in the human pathogen *Listeria monocytogenes*. This route also determines the intrinsic resistance of *L. monocytogenes* against cephalosporins, a group of β-lactam antibiotics. Signaling involves PrkA, a membrane-embedded protein kinase, that is activated during cell wall stress to phosphorylate its target ReoM. Depending on its phosphorylation, ReoM activates or inactivates PG production by controlling the proteolytic stability of MurA, which catalyzes the first step in PG biosynthesis. MurA degradation depends on the ClpCP protease and we here have isolated *murA* mutations that escape this degradation. Using these mutants, we could show that regulation of PG biosynthesis through control of MurA stability is the primary purpose of PrkA-mediated signaling in *L. monocytogenes*. Further experiments identified the transglycosylase RodA and the transpeptidase PBP B3 as additional effectors of PrkA signaling. Our results suggest that both proteins act together to translate the signals received by PrkA into intensification of PG biosynthesis. These findings shed new light on the regulation of PG biosynthesis in Gram-positive bacteria with intrinsic β-lactam resistance.

## Introduction

The cell wall of Gram-positive bacteria is built up from peptidoglycan (PG) strands that are crosslinked with each other to form a three-dimensional network called the sacculus. The sacculus engulfs the entire cell, determines cell shape and provides the key matrix for the attachment of other surface molecules such as teichoic acids, proteins and polymeric carbohydrates. PG biosynthesis is a target of many commonly used antibiotics that can lead to lysis of bacterial cells by weakening the integrity of the sacculus that then gives way to the high internal turgor pressure (1–3). PG accounts for approximately 20% of the weight of a Gram-positive cell (4), illustrating the high demand PG biosynthesis imposes on biosynthetic and energy supplying pathways. Thus, bacteria need to tightly adjust PG production to changing environmental and growth conditions to prevent unnecessary losses of building blocks and energy.

The first step in PG biosynthesis is mediated by MurA, which initiates a series of eight cytoplasmic reactions that sequentially convert UDP-*N*-acetyl-glucosamine (UDP-Glc*N*Ac) into a lipid-linked disaccharide carrying a pentapeptide side chain (lipid II). Lipid II is flipped to the other side of the membrane by Amj- or MurJ-like flippases (5–7), where the disaccharide unit is incorporated into the growing PG chain and the pentapeptides crosslinked either through bifunctional penicillin binding proteins (PBPs) providing transglycosylase and transpeptidase activity (8, 9) or through FtsW/RodA-like transglycosylases that act in concert with monofunctional PBPs, which are mere transpeptidases (10–13).

Among the various regulatory mechanisms controlling PG production, PASTA (<u>P</u>BP <u>a</u>nd <u>s</u>erine/threonine kinase <u>a</u>ssociated) domain-containing <u>e</u>ukaryotic-like <u>s</u>erine/threonine protein <u>k</u>inases (PASTA-eSTKs) have an outstanding role in PG biosynthesis regulation of firmicutes and actinobacteria, two major types of Gram-positives (14–16). PASTA-eSTKs sense cell wall damaging conditions by interaction of lipid II with their extracellular PASTA domains, and this leads to the activation of their cytosolic kinase domain (17, 18). In actinobacteria, many of the kinase substrates directly participate in PG biosynthesis, such as GlmU, important for biosynthesis of UDP-Glc*N*Ac, MviN, a MurJ-like flippase or the bifunctional PBP PonA1 (19–21). Furthermore, actinobacteria control MurA activity by PASTA-eSTK-dependent phosphorylation of CwlM, which allosterically activates MurA (22, 23).

Interestingly, MurA is also the target of PASTA-eSTK-dependent regulation in firmicutes. This involves ReoM, which we have recently described as a novel substrate of the PASTA-eSTK PrkA in *Listeria monocytogenes* (24), a foodborne pathogen causing life-threatening infections (25). We originally identified *reoM* in a screen for suppressors of the heat-sensitive phenotype of a *L. monocytogenes* mutant lacking the late cell division protein GpsB (24, 26, 27). An *L. monocytogenes* Δ*reoM* mutant has increased ceftriaxone resistance and thicker polar PG layers indicating activated PG biosynthesis (24). PrkA phosphorylates ReoM *in vitro* at the conserved Thr-7 residue (24) and PrkA-dependency of this phosphorylation was also demonstrated *in vivo* (28). ReoM is conserved in firmicutes but absent from actinobacteria (24, 29). It does not act as an allosteric activator of MurA but rather controls proteolytic stability of MurA together with ReoY, a second new factor that was identified in the same screen (24). In *L. monocytogenes* and *Bacillus subtilis*, MurA is a substrate of the ClpCP protease (30, 31) and *reoM* and *reoY* mutants accumulate MurA to a similar extent as a *clpC* mutant (24, 27). Analysis of phospho-ablative *reoM* strains and mutants depleted for PrkA and the corresponding protein phosphatase PrpC showed that genetic constellations, in which ReoM is locked in the phosphorylated state, cause MurA stabilization and *vice versa* (24). Thus, PrkA activation leads to activation of PG biosynthesis in firmicutes and actinobacteria even though through different mechanisms.

ReoM interacts with MurA and ReoY in a bacterial two hybrid system and ReoY in turn binds ClpC and ClpP (24). We hypothesize that ReoM and ReoY have an adaptor-like function and present MurA for degradation to the ClpCP protease, but how these proteins exactly contribute to MurA degradation is not known.

We here describe the identification of novel *gpsB* suppressors. The subsequent in-depth analysis of these suppressor mutants led to the identification of MurA variants that escape proteolytic degradation. We use these mutants to demonstrate that control of MurA degradation by ReoM and ClpCP is the major regulatory pathway of PrkA-dependent signaling in *L. monocytogenes* in batch culture as well as during macrophage infection.

## Results

### Novel *gpsB* suppressor mutations in *murA* and *prpC*

An *L. monocytogenes* Δ*gpsB* mutant has a temperature sensitive growth phenotype and cannot multiply at 42°C (26). However, this mutant readily picks up *shg* suppressor mutations (<u>s</u>uppression of <u>h</u>eat-sensitive <u>g</u>rowth) correcting this growth defect and known *shg* suppressor mutations map to *clpC, murZ, reoM* and *reoY*, all involved in regulation of MurA stability by ClpCP-dependent proteolysis (24, 27). In order to identify further mutations suppressing the *gpsB* phenotype, the *L. monocytogenes* Δ*gpsB* mutant strain LMJR19 was streaked on BHI plates and incubated at 42°C. After two days of incubation, 50 suppressors were isolated and streaked to single colonies. The *clpC, murZ, reoM* and *reoY* genes of these 50 suppressors were amplified by PCR and sequenced by Sanger sequencing. Seven suppressors had wild type alleles of the four known *gpsB* suppressor genes and thus must have acquired novel *shg* suppressor mutations somewhere else on the chromosome. Genome sequencing of these seven *shg* suppressors revealed unique mutations mapping either to *murA* or to *prpC*. While the *murA* gene was affected by mutations introducing amino acid substitutions only (*shg19*: *murA S262L, shg21*: *murA N197D*), the *shg* mutations in *prpC* introduced amino acid exchanges (*shg24*: *prpC N83S L125F, shg47*: *prpC G39S*) as well as premature stop codons (*shg32*: *prpC*^*1-109*^, *shg42*: *prpC*^*1-73*^, *shg55*: *prpC*^*1-44*^). This observation is in good agreement with the reported essentiality of *murA* (27), but conflicts with our previous observation that *prpC* could not be removed from the *L. monocytogenes* genome (24). Suppressor strains *shg19* (Δ*gpsB murA S262L*), *shg21* (Δ*gpsB murA N197D*) and *shg55* (Δ*gpsB prpC*^*1-44*^) were chosen for further experiments and growth was recorded at 37°C and 42°C. All three suppressor strains grew almost like wild type at 37°C and 42°C, whereas the Δ*gpsB* mutant grew with delayed rate at 37°C and did not grow at 42°C (Fig 1A-B).

**Figure 1:**
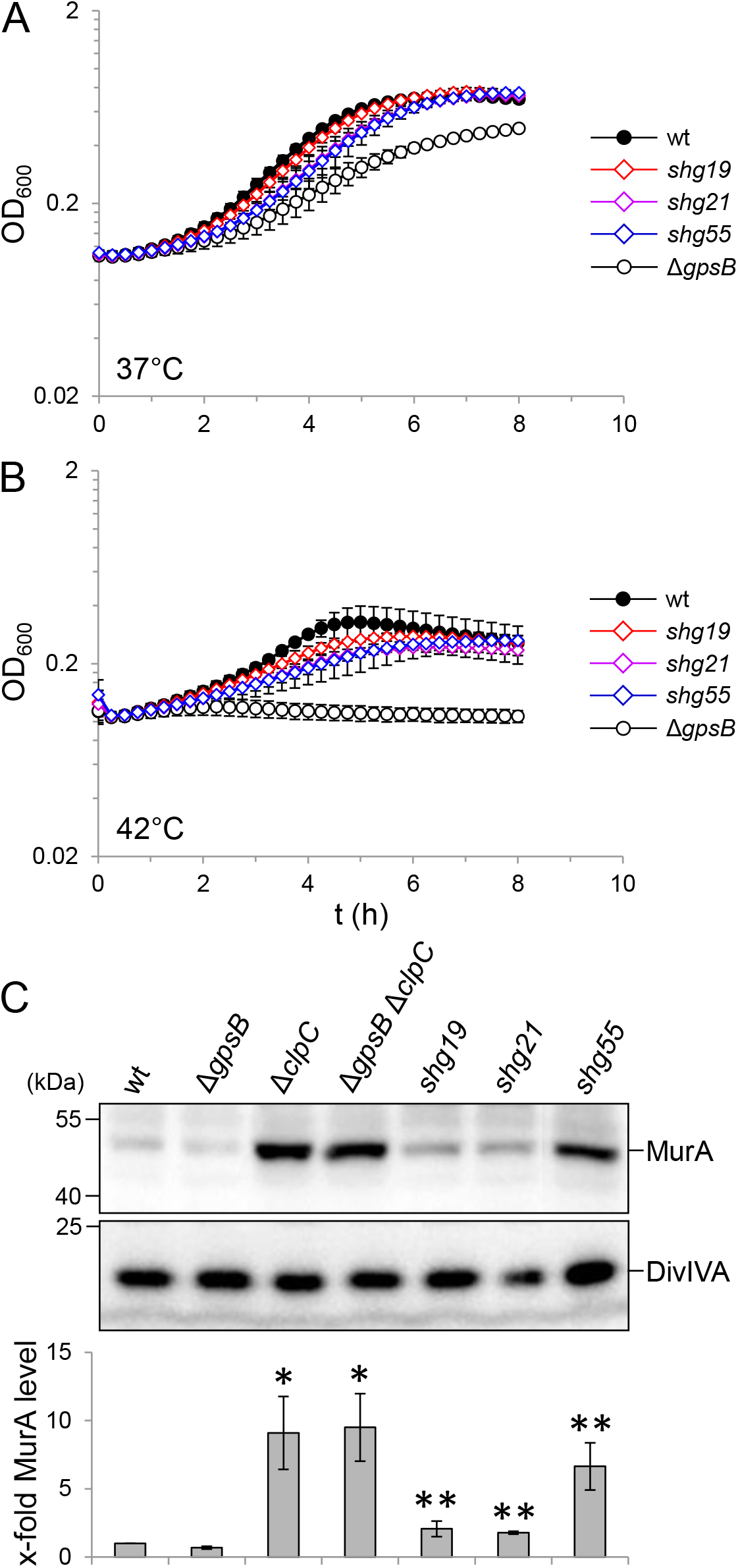
Suppression of the Δ*gpsB* growth defect by *murA* and *prpC* mutations. (A-B) Growth of *L. monocytogenes* strains with mutations in *gpsB, murA* and *prpC. L. monocytogenes* strains EGD-e (wt), LMJR19 (Δ*gpsB*), *shg19* (Δ*gpsB murA S262L*), *shg21* (Δ*gpsB murA N197D*) and *shg55* (Δ*gpsB prpC*^*1-44*^) were grown in BHI broth at 37°C (A) and 42°C (B). The experiment was performed in triplicate and average values and standard deviations are shown. (C) Western blot showing MurA and DivIVA levels (for control) in the same set of strains. Strains LMJR138 (Δ*clpC*) and LMJR139 (Δ*gpsB* Δ*clpC*) were included as controls. Equal amounts of total cellular proteins were loaded. MurA signals from three independent experiments were quantified by densitometry and average values and standard deviations are shown. Asterisks indicated statistically significant differences compared to wild type (*) or compared to the Δ*gpsB* mutant (**, *t*-test, *P*<0.05).

Suppression in *shg* suppressors can involve proteolytic stabilization of MurA leading to stimulation of PG biosynthesis. In order to test, whether MurA is stabilized in the novel *shg* suppressors, their cellular MurA levels were determined by Western blotting. This revealed a mild increase of the MurA level in *shg19* (2.1±0.6-fold compared to wild type) and *shg21* (1.8±0.1-fold) and a strong increase in *shg55* (6.6±1.7-fold), which is almost as strong as observed in a Δ*clpC* mutant (9.1±2.7-fold, Fig 1C). Taken together, we have identified novel *shg* suppressor mutations in *murA* and *prpC* affecting the cellular levels of MurA.

### Deletion of *prpC* is tolerated in the Δ*gpsB* background

Previous results demonstrated that *prpC* could not be removed from the chromosome unless compensatory *prkA* mutations reduced the activity of the cognate protein kinase (24). As this suggested that *prpC* represents an essential gene in *L. monocytogenes*, we were surprised to see that *prpC* repeatedly was inactivated by the introduction of premature stop codons in the *gpsB* suppressor mutants *shg32, shg42* and *shg55*. To test whether *prpC* becomes dispensable in the absence of *gpsB*, we tried to delete *prpC* in the background of the Δ*gpsB* mutant using integration/re-excision of a temperature sensitive plasmid designed to remove the *prpC* gene during excision from the chromosome. Plasmid excision is supposed to generate a 50:50 mixture of wild type and deletion mutant clones (32). Unlike in previous *prpC* deletion attempts in wild type background (24), *prpC* could be readily deleted in the Δ*gpsB* mutant. Furthermore, it even seemed that *prpC* deletion is favored in the absence of *gpsB*, as 19 out of 20 tested clones had factually lost *prpC*. Unlike the Δ*gpsB* single mutant, the resulting Δ*gpsB* Δ*prpC* double mutant could grow at 42°C, even though it did not fully reached normal wild type growth (Fig S1A). Moreover, MurA levels increased 2.8±0.5-fold in the Δ*gpsB* Δ*prpC* double mutant compared to wild type (Fig S1B). These data confirm that inactivation of *prpC* suppresses the Δ*gpsB* phenotype and that suppression involves accumulation of MurA.

### Effect of N197D and S262L mutations on MurA activity

MurA proteins consist of two globular domains connected to each other. The UDP-Glc*N*Ac binding site and the catalytic center are located at the interface between these two domains (33). The MurA residues replaced in the two *shg* suppressors are located outside the active site in helical regions exposed at the surface of the N-terminal (N197D) and the C-terminal (S262L) domain (Fig S2A). We assumed that these mutations would improve MurA activity to exert a similar suppressing effect on the Δ*gpsB* mutant as increased MurA levels. To test this, we generated strains that carry IPTG-inducible *murA* genes (wild type *murA* as well as S262L and N197D variants) at the *attB* site but have their endogenous *murA* genes removed. Consistent with the essentiality of *murA* (27), all three strains required IPTG for growth in BHI broth (Fig S2B), when their pre-cultures were cultivated in the absence of IPTG. However, growth of strains LMSW140 (i*murA S262L*) and LMSW141 (i*murA N197D*) was similar to strain LMJR123 (i*murA*) in the presence of 1 mM IPTG (Fig S2B) and also at lower IPTG concentrations (data not shown). Next, we measured the susceptibility of these strains to fosfomycin, a known inhibitor of MurA (34). Fosfomycin sensitivity was measured in a disc diffusion assay on BHI agar plates, which allowed growth of the inducible *murA* strains even in the absence of IPTG due to background *murA* expression. Sensitivity of these strains towards fosfomycin was increased compared to wild type due to MurA depletion, but effects of the N197D and S262L mutations did not become evident (Fig S2C).

To further test the possibility that the two *murA* mutations altered enzyme activity, wild type MurA, MurA N197D and MurA S262L were purified as Strep-tagged proteins to near homogeneity (Fig S2D) and their enzymatic activity was tested. This showed that the MurA N197D protein was as active as wild type MurA, whereas the activity of MurA S262L was reduced (Fig S2E). Taken together, these results rule out the possibility that increased MurA activities explain suppression of the Δ*gpsB* phenotype.

### The MurA N197D and S262L variants escape proteolytic degradation

Next, we wondered whether the two *murA* mutations affect proteolytic degradation of MurA. In order to test this idea, the *gpsB* deletion in the two suppressor strains *shg19* (Δ*gpsB murA S262L*) and *shg21* (Δ*gpsB murA N197D*) was repaired by re-introduction of the native *gpsB* allele at the original locus to generate strains that carry the two *murA* mutations as the sole genetic changes in comparison to wild type. MurA degradation in the resulting strains LMSW155 (*murA S262L*) and LMSW156 (*murA N197D*) was then compared to wild type and the Δ*clpC* mutant. For this purpose, these strains were grown to an OD_600_ of 1.0 and 100 µg/ml chloramphenicol was added to block protein biosynthesis. MurA levels over time were then determined by Western blotting. As reported previously (24), MurA levels rapidly declined over time in wild type cells, but stayed stable in a Δ*clpC* mutant (Fig 2A-B), reflecting ClpCP-dependent proteolytic degradation of MurA. However, degradation of both MurA mutant proteins was significantly delayed (Fig 2A-B), which is in good agreement with their increased levels in the original *shg19* and *shg21* suppressors (Fig 1C). This shows that the MurA N197D and S262L variants escape proteolytic degradation by ClpCP.

**Figure 2:**
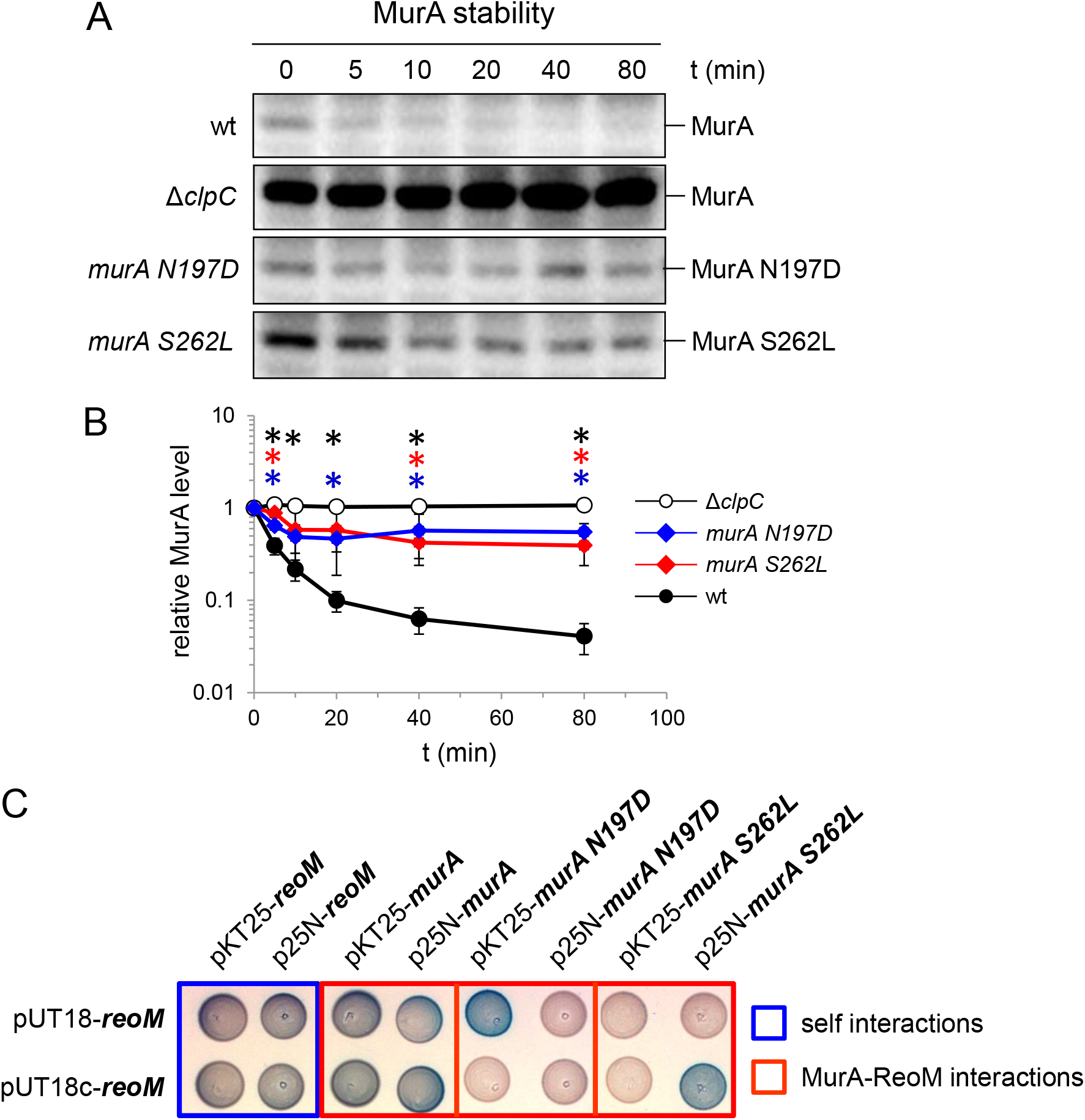
Effect of the N197D and S262L substitutions on MurA stability. (A) Western blots following MurA protein degradation *in vivo. L. monocytogenes* strains EGD-e (wt), LMJR138 (Δ*clpC*), LMSW155 (*murA S262L*) and LMSW156 (*murA N197D*) were cultivated to an OD_600_ of 1.0 and 100 µg/ml chloramphenicol was added to stop protein biosynthesis. Samples were taken before chloramphenicol addition and after different time points to analyze MurA levels. (B) MurA signals were quantified using ImageJ and average values and standard deviations are shown (n=3). MurA levels of each strain before chloramphenicol addition (t = 0 min) were arbitrarily set to 1 and the remaining signal intensities at later time points are shown as relative values. Statistically significant differences (compared to wild type) are marked with asterisks (*P*<0.05, *t*-test). (C) Bacterial two hybrid assay to test the interaction of ReoM with MurA and its N197D and S262L variants.

Previously published bacterial two hybrid data indicated that MurA directly binds ReoM, which – together with ReoY – could present MurA to ClpCP (24). We wondered whether the MurA N197D and S262L mutants have lost the ability to interact with ReoM and tested this in the bacterial two hybrid system. As published before (24), B2H detects an interaction of MurA with ReoM. However, the MurA N197D and S262L proteins have each lost their interaction with MurA in several of the four possible permutations (Fig 2C), even though they still interacted with itself (data not shown). This raises the interesting possibility that the two MurA variants could be stabilized because they do not longer bind to ReoM.

### Escape mutations in *murA* affect peptidoglycan biosynthesis and growth

MurA was shown to be a key switch in regulation of PG biosynthesis and MurA levels strictly correlated with PG production and ceftriaxone resistance in *L. monocytogenes* (24, 27). Recent work in *B. subtilis* further confirmed this central role of MurA in PG biosynthesis regulation (35). In order to test whether the two *murA* escape mutations influence PG production, their ceftriaxone resistance was tested and the Δ*clpC* and the i*murA* mutants were included as control. Ceftriaxone resistance was more than ten-fold increased in the Δ*clpC* mutant and inducer-dependent in the IPTG-controlled i*murA* strain as shown previously (24). However, ceftriaxone resistance was also 3-4-fold higher in the two *murA* escape mutants, indicating that stabilization of MurA in these two strains indeed caused stimulation of PG biosynthesis (Fig 3A). Fluorescence microscopy of nile red-stained cells also revealed changes in cellular morphology, as cells of the two *murA* escape and Δ*clpC* mutants were clearly thinner (Fig 3B), whereas cell length was not affected (data not shown). Apparently, MurA stabilization also impairs maintenance of normal cell width.

**Figure 3:**
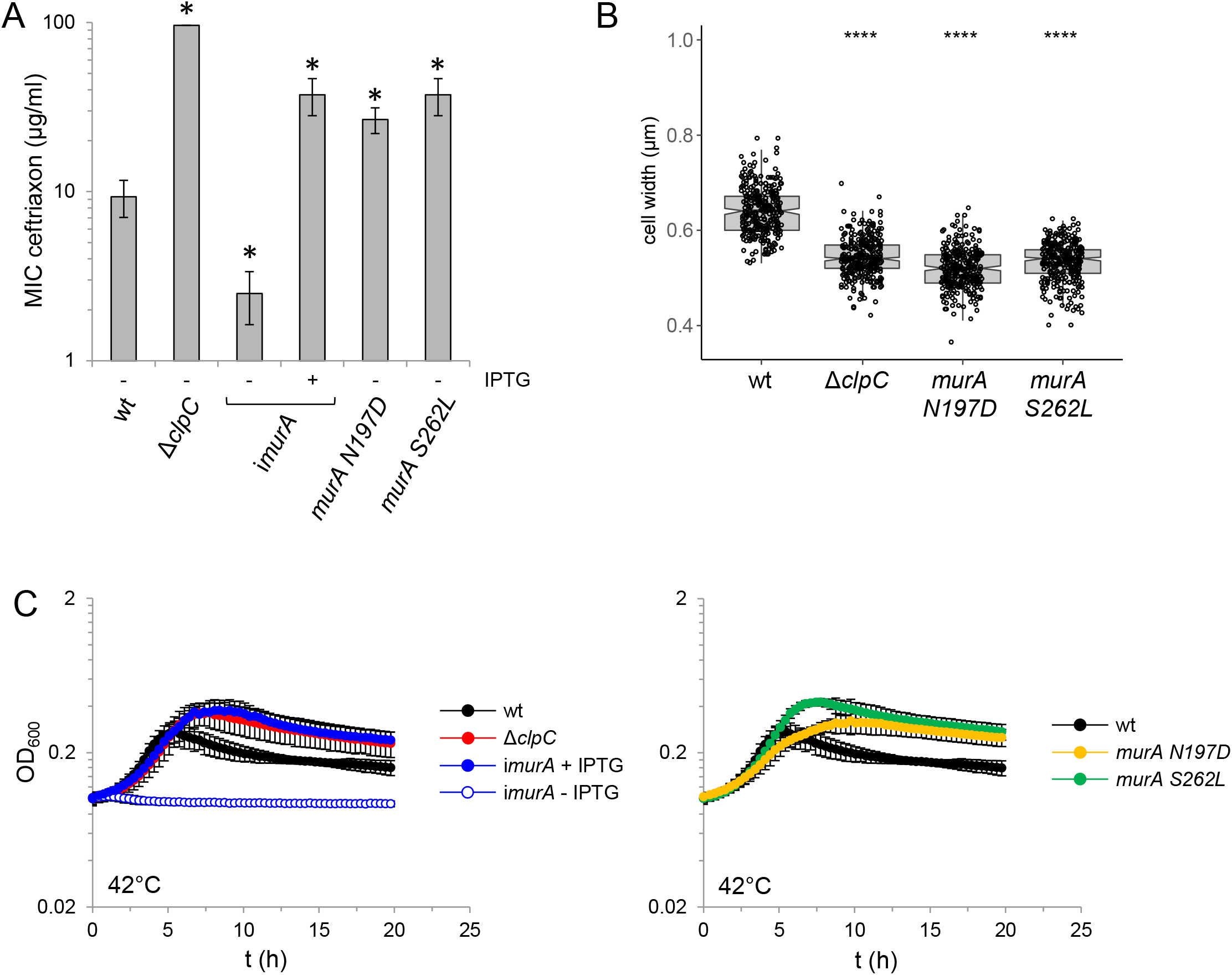
Effect of *murA* escape mutations on ceftriaxone resistance, morphology and growth at high temperature. (A) Minimal inhibitory concentration (MIC) of ceftriaxone of *L. monocytogenes* strains EGD-e (wt), LMJR138 (Δ*clpC*), LMSW155 (*murA S262L*) and LMSW156 (*murA N197D*) as determined by E-tests. Strain LMJR123 (i*murA*) grown in the absence or presence of 1 mM IPTG was included for control. Average values and standard deviations are shown (n=3). Asterisks mark statistically significant differences (*t*-test, *P*<0.01). (B) Cell diameters of *L. monocytogenes* strains EGD-e (wt), LMJR138 (Δ*clpC*), LMSW155 (*murA S262L*) and LMSW156 (*murA N197D*) during logarithmic growth in BHI broth at 37°C. Cells were stained with nile red and cell width of 300 cells per strain was measured. The experiment was repeated three times and one representative experiment is shown. Asterisks indicate significance levels (**** - *P*<0.0001, *t*-test). (C) Growth of the same set of strains as in panel A in BHI broth at 42°C. For clarity, growth curves were split into two diagrams. Average values and standard deviations from an experiment performed in triplicate are shown.

Differences in growth of the *murA* escape mutants were not detected in BHI broth at 37°C, but a characteristic pattern that correlates well with the anticipated MurA expression levels became evident at 42°C. Here, the Δ*clpC* mutant grew to a higher optical density in stationary phase and a congruent growth curve was observed with the i*murA* strain in the presence of IPTG (Fig 3C). We have shown earlier that both strains highly overexpress MurA under these conditions (27). Remarkably, both *murA* escape mutants also lead to higher stationary phase optical densities at 42°C (Fig 3C), suggesting that proteolytic degradation of MurA becomes important at higher temperature to limit bacterial growth. Analysis of the ultrastructure of the PG layer under these conditions revealed thicker PG layers in *murA N197D, murA S262L* and Δ*clpC* mutants, particularly at the poles (Fig S3A-B). Moreover, polar PG thickness was found to be IPTG-dependent in a strain overexpressing *murA* (Fig S3A-B). This unequivocally shows that control of PG production through proteolytic stabilization of MurA is a key switch to control listerial growth and PG ultrastructure, at least at higher temperature.

That the two *murA* escape mutations with their positive effects on growth and antibiotic resistance have not succeeded during evolution suggests that they are also associated with evolutionary disadvantages. Accordingly, the *murA N197D* and *S262L* mutants were found to grow slower in the presence of salt (Fig S4A) and were more sensitive against lysozyme (Fig S4B). This shows that increased biomass yields that are caused by MurA overproduction comes at the price of increased sensitivity against different other stressors.

### The *murA N197D* escape mutation uncouples viability from PrkA signaling

The *prkA* gene is essential because phosphorylated ReoM keeps ClpCP in control to prevent unregulated proteolysis of its essential substrate MurA (24). Following with this model, MurA variants that escape proteolytic degradation through ClpCP should release the cell from PrkA dependency since ClpCP (even when overactive in the absence of PrkA and phosphorylated ReoM) does not longer accept such MurA variants as substrates. In order to test this hypothesis, we tried to delete *prkA* in *murA N197D* and *murA S262L* backgrounds, which never has been possible in the wild type background of *L. monocytogenes* EGD-e (24). Similarly, the *prkA* gene could not be removed in the *murA S262L* background, presumably because of the reduced activity of MurA S262L. In contrast, *prkA* could be easily deleted in the *murA N197D* strain. The resulting Δ*prkA murA N197D* double mutant was viable and grew like the wild type, whereas PrkA-depleted cells could not grow at all (Fig 4A). Moreover, the Δ*prkA murA N197D* mutant was viable inside macrophages and even grew intercellularly with a wild type-like growth rate. This was in complete contrast to PrkA-depleted cells (Fig 4B), suggesting that even the intracellular growth defect upon PrkA depletion, may entirely be MurA-dependent. Ultimately, the extreme ceftriaxone sensitivity seen in PrkA-depleted cells did not develop in the Δ*prkA murA N197D* strain (Fig 4C). These results demonstrate that PrkA becomes non-essential for viability, virulence and PG biosynthesis in mutants, in which MurA escapes proteolytic degradation by ClpCP. These data collectively indicate that MurA is the relevant target of PrkA signaling in *L. monocytogenes* during standard growth conditions and during growth inside macrophages. This extends previous conclusions on the relevance of ReoM for PrkA signalling (24, 28) as it suggests that ReoM is specific for MurA and likely does not control the degradation of additional essential substrates in *L. monocytogenes*. In agreement with this, deletion of the *reoM, reoY* and *murZ* genes, which are all involved in control of MurA degradation, suppressed lethality and ceftriaxone susceptibility of the Δ*prkA* deletion – albeit to different degrees (Fig S5).

**Figure 4:**
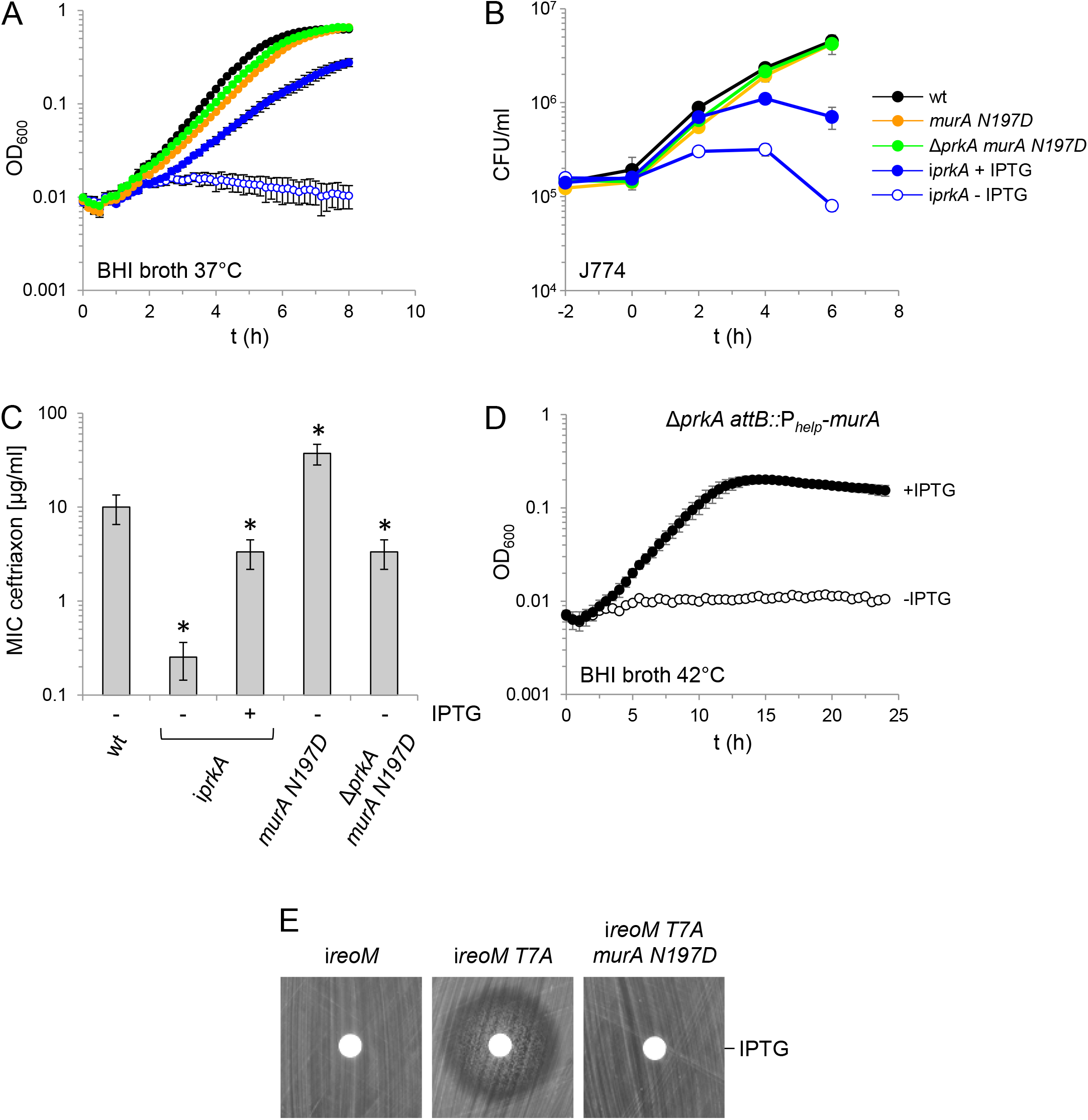
Suppression of lethal *prkA* and *reoM* phenotypes by *murA* mutations. (A) Growth of *L. monocytogenes* strains EGD-e (wt), LMSW84 (i*prkA*), LMSW156 (*murA N197D*) and LMS266 (Δ*prkA murA N197D*) in BHI broth at 37°C. Average values and standard deviations from technical replicates (n=3) are shown. (B) Intracellular growth of strains EGD-e (wt), LMSW156 (*murA N197D*) and LMS266 (Δ*prkA murA N197D*) in J774 macrophages. The experiment was performed as triplicate and average values and standard deviations are shown. (C) Minimal inhibitory ceftriaxone concentrations of the same set of strains as in panels A and B. Asterisks mark statistically significant differences compared to wild type (n=3, *P*<0.05, *t*-test). (D) MurA overexpression rescues the Δ*prkA* mutant. Growth of strains LMS272 (Δ*prkA attB::*P_*help*_*-murA*) in BHI broth ± 1 mM IPTG at 42°C. The experiment was carried out as triplicate and average values and standard deviations are shown. (E) The *murA N197D* mutation suppresses *reoM T7A* toxicity. Disc diffusion assay with filter discs loaded with 10 µl of a 1 M IPTG solution. Strains used were LMSW57 (i*reoM*), LMSW52 (i*reoM T7A*) and LMS273 (i*reoM T7A murA N197D*).

To further substantiate these claims, we tested whether overexpression of MurA would also rescue Δ*prkA* lethality. As hypothesized by this idea, deletion of *prkA*, which is impossible in wild type (24, 36), turned out to be possible in strain LMJR116, in which a second copy of *murA* was expressed from an IPTG-dependent promoter. Growth of the resulting strain (LMS272) was not IPTG-dependent at 37°C (data not shown), probably due to leakiness of the IPTG-controlled promoter driving a certain degree of expression of the second *murA* copy even in the absence of IPTG. However, IPTG was required to support growth of this strain at 42°C (Fig 4D). The *clpC* gene is strongly induced at 42°C (37), leading to stimulated MurA degradation. This explains why IPTG-dependency of this strain was temperature dependent. However, as the main outcome, this experiment clearly demonstrated that overexpression of MurA is sufficient to suppress the essentiality of *prkA*, which is in accordance with the idea that control of MurA stability is the main purpose of PrkA signaling.

We have shown in previous work that expression of *reoM T7A* is toxic to *L. monocytogenes* because ClpCP is overactive when ReoM cannot be phosphorylated at Thr-7 by PrkA (24). If MurA is indeed the main subject of PrkA signaling, then the *murA N197D* mutation should also overcome the toxicity of the phospho-ablative *reoM T7A* variant. To test this idea, a *murA N197D* strain was generated, in which expression of *reoM T7A* can be controlled by IPTG (strain LMS273) and susceptibility of this strain against IPTG was tested in a disc diffusion assay. While IPTG was clearly toxic for strain LMSW52 (i*reoM T7A*), strain LMS273 fully tolerated IPTG (Fig 4E). This further confirms that the main role of the PrkA:ReoM signaling axis is control of MurA stability.

### ClpC-dependent activation of PG biosynthesis requires PBP B3 and RodA3

Many lines of evidence congruently show that stabilization of MurA enhances PG formation and that ceftriaxone sensitivity can be used as a proxy to detect stimulating or disadvantageous effects on PG biosynthesis (this work) (24, 27). While most enzymes necessary for the cytoplasmic steps of PG biosynthesis are non-redundant, there is a high degree of redundancy in *L. monocytogenes* genes encoding PBPs and FtsW/RodA-like transglycosylases, required for the final steps outside the cell (38, 39). Inactivation of the class A PBPs had only mild effects on cephalosporin resistance in previous experiments (40), and were not further considered here. However, PBP B3, one out of three class B PBPs, substantially contributed to cephalosporine resistance (40, 41). Likewise, deletion of *rodA1* and *rodA2* only slightly affected cephalosporin resistance, whereas deletion of *rodA3* had a greater impact (39). Thus, it seemed likely that PBP B3 and RodA3 cooperate to ensure activation of PG biosynthesis when MurA accumulates.

In order to test this hypothesis, we tested the effect of *pbpB3* and *rodA3* mutations on the ceftriaxone resistance level of the Δ*clpC* mutant (MIC 96±0 µg/ml), which is strongly increased compared to wildtype (9.3±2.3 µg/ml, Fig 5A). As deletion of *pbpB3* was not possible in the Δ*clpC* background (data not shown), *clpC* was deleted in an IPTG-inducible i*pbpB3* mutant. Depletion of PBP B3 in the i*pbpB3* mutant caused a similar reduction in ceftriaxone resistance (MIC: 0.8±0 µg/ml) as observed in a Δ*pbpB3* strain (MIC: 0.6±0.1, Fig 5A). Remarkably, reduction of ceftriaxone resistance almost down to the level of a Δ*pbpB3* mutant was also observed in the i*pbpB3* Δ*clpC* strain, when grown in the absence of IPTG (1.0±0 µg/ml). This demonstrates that the increased ceftriaxone resistance of the Δ*clpC* mutant is PBP B3-dependent. Next, the *rodA2-rodA1* gene pair and the *rodA3* gene were inactivated in the Δ*clpC* mutant. While ceftriaxone resistance was only slightly affected by the Δ*rodA2-rodA1* deletion (MIC 6.7±1.2 µg/ml) compared to wildtype (10.7±2.4 µg/ml in this experiment), inactivation of *rodA3* had a somewhat stronger effect (5.3±1.2 µg/ml). More interestingly, the increased ceftriaxone resistance level of the Δ*clpC* mutant (96±0 µg/ml) was not affected by the Δ*rodA2-rodA1* deletion (74.7±18.5 µg/ml), but was reduced back almost to wildtype level when *rodA3* was inactivated (13.3±2.3 µg/ml, Fig 5B). All in all, this shows that PBP B3 and RodA3 together form a cognate PBP-FtsW/RodA pair. Furthermore, this couple is required for PG biosynthesis activation as it occurs under conditions leading to proteolytic stabilization of MurA.

**Figure 5:**
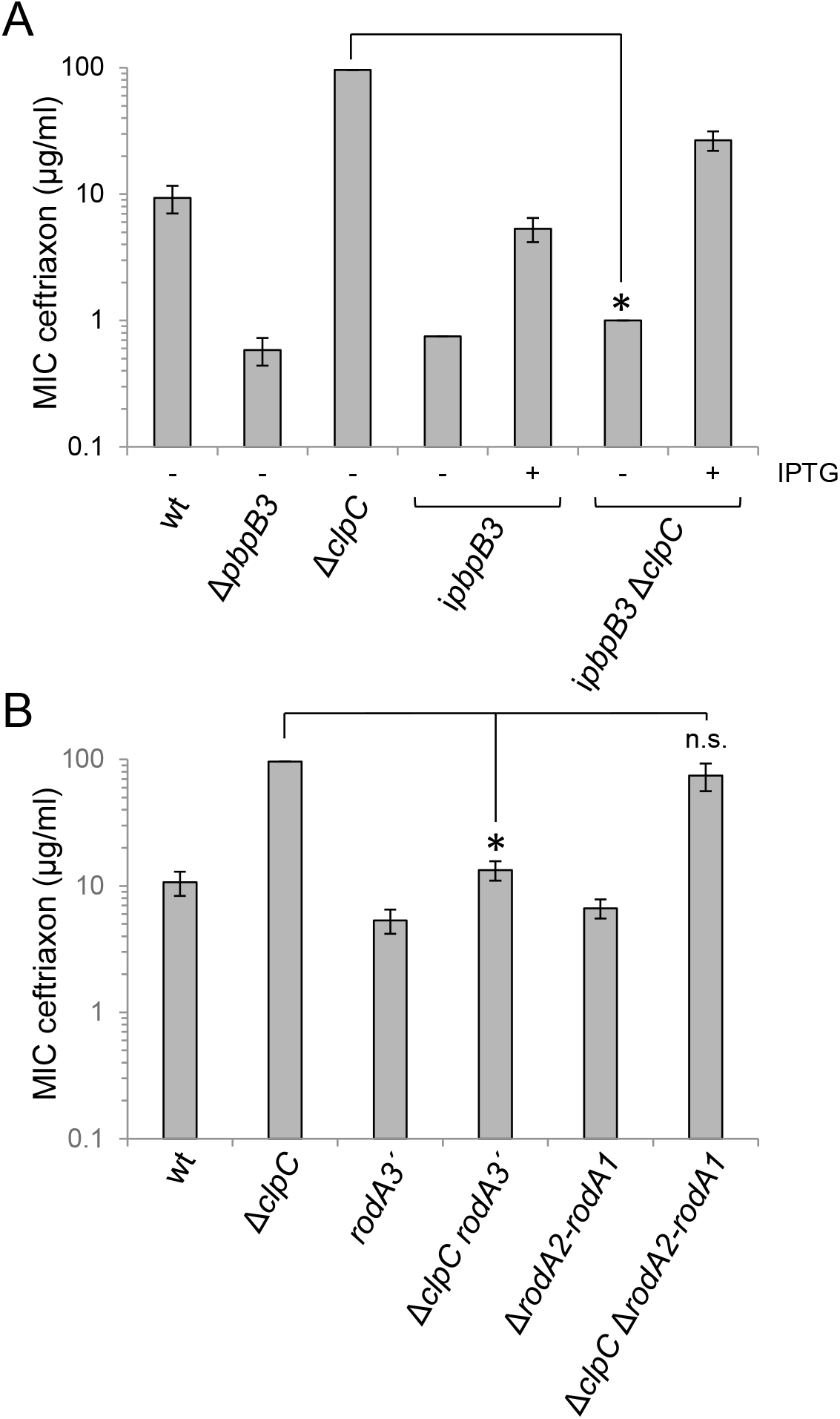
ClpC-dependent activation of PG biosynthesis requires RodA3 and PBP B3. (A) Depletion of PBP B3 suppressed the increased ceftriaxone resistance of the Δ*clpC* mutant. Minimal inhibitory ceftriaxone concentrations of *L. monocytogenes* strains EGD-e (wt), LMJR41 (Δ*pbpB3*), LMJR138 (Δ*clpC*), LMMF1 (i*pbpB3*) and LMSW143 (i*pbpB3* Δ*clpC*) grown on agar plates ± 1 mM IPTG. (B) Inactivation of *rodA3* suppresses the increased ceftriaxone resistance of the Δ*clpC* mutant. Minimal inhibitory ceftriaxone concentrations of *L. monocytogenes* strains EGD-e (wt), LMJR138 (Δ*clpC*), LMS267 (*rodA3’*), LMS268 (Δ*clpC rodA3’*), LMSH67 (Δ*rodA2-rodA1*) and LMS270 (Δ*clpC* Δ*rodA2-rodA1*). MICs were determined using E-tests and average values and standard deviations calculated from three repetitions are shown. Asterisks indicate statistically significant differences (*t*-test, *P*<0.01).

## Discussion

Many substrates of PASTA-eSTKs have been identified in the different Gram-positive bacteria. Among these kinase substrates are proteins from various cellular pathways, but some functional classes are overrepresented: (i) Proteins acting in carbon and cell wall metabolism such as the HPr and YvcK proteins of *B. subtilis* and *L. monocytogenes* (28, 42–44), (ii) regulatory proteins such as the WalR and GraR response regulators of *B. subtilis* and *Staphylococcus aureus*, respectively (45, 46), and (iii) proteins acting in cell division, among which GpsB is one of the proteins consistently found as a PASTA-eSTK substrate in *B. subtilis, L. monocytogenes* and *Streptococcus pneumoniae* (28, 47, 48). This diversity of substrates and their own – often pleiotropic - functions have impeded the identification of primary and the discrimination of less important kinase substrates. We previously have identified ReoM as a substrate of PrkA in *L. monocytogenes* reference strain EGD-e and could show that deletion of *reoM* suppresses loss of viability that results from PrkA depletion (24). Moreover, expression of a phospho-ablative *reoM T7A* allele was toxic on its own and thus phenocopied the absence of PrkA (24). These tight geno- and phenotypic relations between the PrkA kinase and ReoM, which is only one out of 23 known PrkA substrates in *L. monocytogenes* (28), suggested that ReoM must be a relevant kinase target. We here further support this conclusion by our observation that *prkA* even can be deleted in a Δ*reoM* background, again indicating that the phosphorylation of ReoM is the particular PrkA-dependent phosphorylation event that is crucial for viability. Remarkably, Kelliher *et al*. reported that *prkA* is non-essential in *L. monocytogenes* strain 10403S (28) and discuss strain-specific differences as an explanation. In fact, *L. monocytogenes* strains EGD-e and 10403S are relatively distinct with respect to the population structure of the species *L. monocytogenes* and even belong to different molecular serogroups (EGD-e: IIc, 10403S: IIa) (49). Thus, differences in the stringency of MurA degradation by ClpCP, in expression of the genes acting in the ReoM/ClpCP/MurA axis or even in the relative expression of *murA* and its paralogue *murZ* may account for this difference. Despite this discrepancy and consistent with our findings, *reoM* deletion suppressed all *prkA* associated phenotypes also in strain 10403S (28).

How ReoM stimulates MurA degradation is presently not known, but it may act as a ClpCP activator or as an adaptor, which presents substrates to the protease complex. ReoM could either be specific for MurA or could also control the stability of other proteins. Our observation that MurA variants exist that escape proteolytic degradation, further support the idea that MurA is a protease substrate even though this has never been confirmed *in vitro* (24, 27, 30). Moreover, the observations that their amino acid substitutions are located on the MurA surface and impair the interaction with ReoM would be consistent with the adaptor hypothesis.

A central result of our study is the finding that one of the *murA* escape mutations (N197D) suppressed *prkA* essentiality under standard growth conditions and during intracellular growth in macrophages, to which *prkA* mutants are specifically sensitive (44). This particular *murA* mutation does not alter enzymatic activity and only rescues MurA from proteolytic degradation. It also overcomes the lethality of the *reoM T7A* allele, the expression of which normally would lead to rapid MurA degradation. These findings provide a genetic answer to the question whether other *L. monocytogenes* proteins exist, the stability of which depends on ReoM phosphorylation: As uncoupling of MurA from PrkA-dependent control of ReoM-mediated ClpCP activation restores otherwise lethal *prkA* and *reoM* phenotypes, we have to conclude that MurA is not only the main substrate or target of ReoM, but also that control of MurA stability is the main purpose of PrkA-mediated signaling in *L. monocytogenes*. This conclusion is further substantiated by the observation that artificial MurA overexpression also rescues the lethal Δ*prkA* phenotype. According to our present model, PrkA gets activated during cell wall stress and phosphorylation of ReoM limits MurA degradation (Fig 6). PG biosynthesis is activated when MurA accumulates. How PG biosynthesis activation under these conditions alters the chemical structure of the sacculus is not known, but a thicker cell wall is produced, and this occurs concomitantly with an increase of ceftriaxone resistance. As a second important result of our study we could show that this latter effect depends on RodA3 and PBP B3, which apparently constitute a cognate glycosyltransferase/transpeptidase pair required during this special mode of PG biosynthesis (Fig 6). In our current model, the PrkA→ReoM/ReoY→MurA/ClpCP→RodA3/PBP B3 cascade would represent a closed homeostatic circuit that senses lipid II (by PrkA) to control its production (through control of MurA stability) and consumption (by RodA3/PBP B3) (Fig 6). The genes in this pathway collectively determine the intrinsic resistance of *L. monocytogenes* to cephalosporins (24, 36, 39, 40, 50) and are conserved in several other Gram-positive bacteria (Fig S6) In *Enterococcus faecalis*, most of the corresponding homologues were shown to maintain the high intrinsic cephalosporin resistance level of this organism (29, 51–54), and deletion of *pbpC*, which encodes the PBP B3 homologue of *B. subtilis* (Fig S6), was also necessary for cephalosporin resistance (55). Moreover, inactivation of Stk1, the PASTA-eSTK of *S. aureus*, sensitizes methicillin resistant *S. aureus* (MRSA) against β-lactams including oxacillin and this is overcome by *reoM* and *reoY* deletions (28, 56). Interestingly, MecA, which represents the major methicillin resistance determinant of MRSA (57), is the equivalent homologue of *L. monocytogenes* PBP B3 (Fig S6) (8). Lastly, the PrkA signaling cascade lacks ReoY and a homologue of PBP B3 in *S. pneumoniae*, which is typically susceptible to penicillins. This raises the interesting possibility that this cascade generally supports a specific type of PG biosynthesis which confers a higher resistance against cephalosporins and other β-lactams. Further studies, also including other bacterial species, would have to be performed to test this hypothesis.

**Figure 6:**
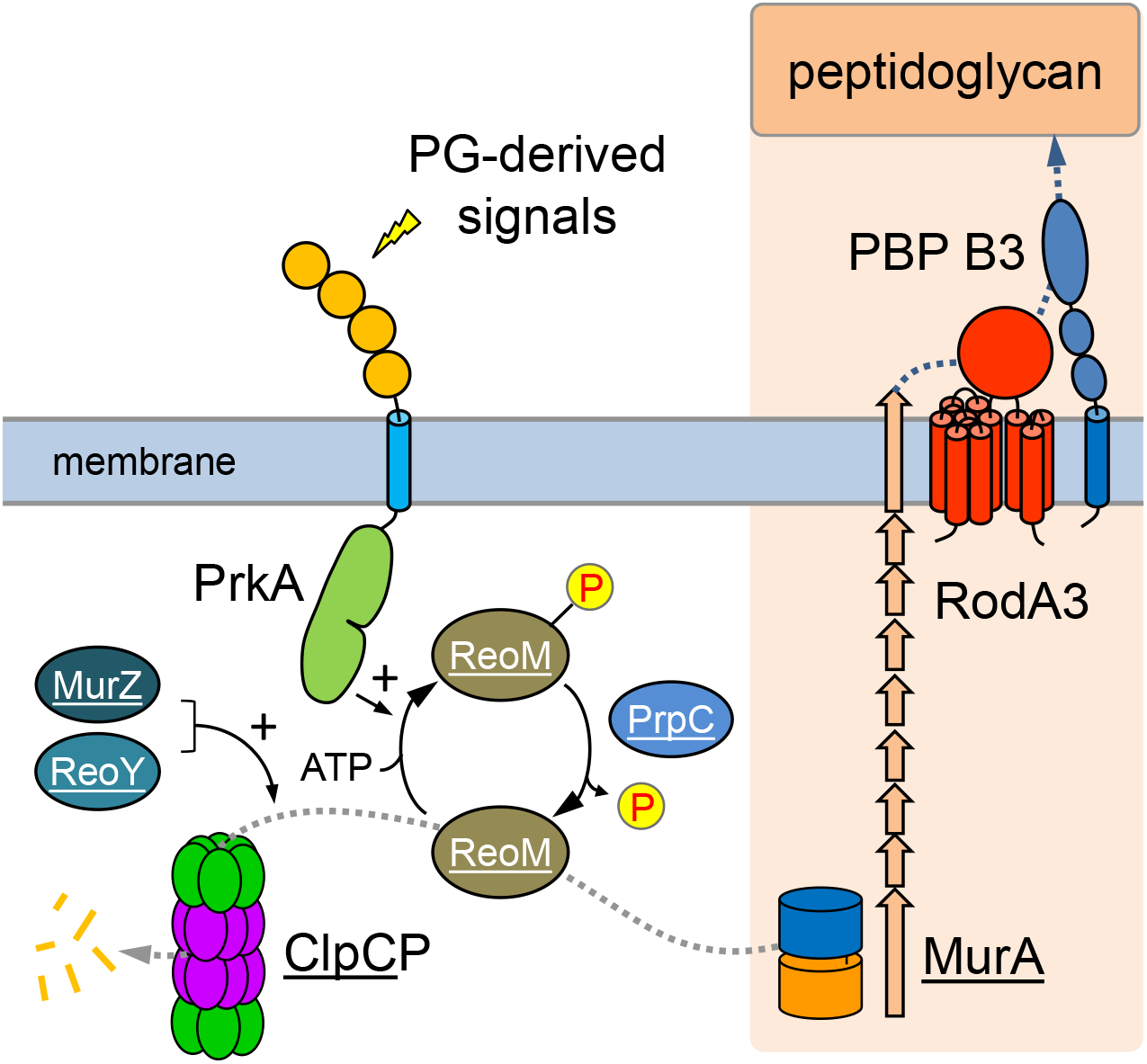
Activation of *L. monocytogenes* PG biosynthesis through PrkA signaling. PrkA phosphorylates ReoM upon activation by PG-derived signals such as lipid II. P-ReoM no longer supports/activates ClpCP-dependent MurA degradation so that MurA can accumulate and PG biosynthesis can take place. Activation of PG biosynthesis under conditions leading to PrkA activation and MurA accumulation requires RodA3 and PBP B3. Underlined proteins have been identified as suppressors of the heat sensitive Δ*gpsB* mutant, which is affected in PG biosynthesis, in this and previous work (24, 27). Image with modifications from (69).

## Materials and Methods

### Bacterial strains and growth conditions

Tab 1 lists all strains used in this study. *L. monocytogenes* strains were routinely cultivated in brain heart infusion (BHI) broth or on BHI agar plates. Antibiotics and supplements were added when required at the following concentrations: erythromycin (5 µg/ml), kanamycin (50 µg/ml), X-Gal (100 µg/ml) and IPTG (as indicated). *Escherichia coli* TOP10 was used as standard host for all cloning procedures (58). Minimal inhibitory concentrations against ceftriaxone were determined as described previously using E-test strips with a ceftriaxone concentration range of 0.016 - 256 µg/ml (Bestbion^dx^) (38). Susceptibility against fosfomycin was tested in a disc diffusion assay using filter discs loaded with 8 µl of a 20 mg/ml fosfomycin solution. For this purpose, *L. monocytogenes* colonies, grown on BHI agar plates, were resuspended in BHI broth and used to swab-inoculate BHI agar plates. Filter discs were spotted on BHI agar plates and incubated over night at 37°C to measure the inhibition zone diameter the next day.

**Table 1:**
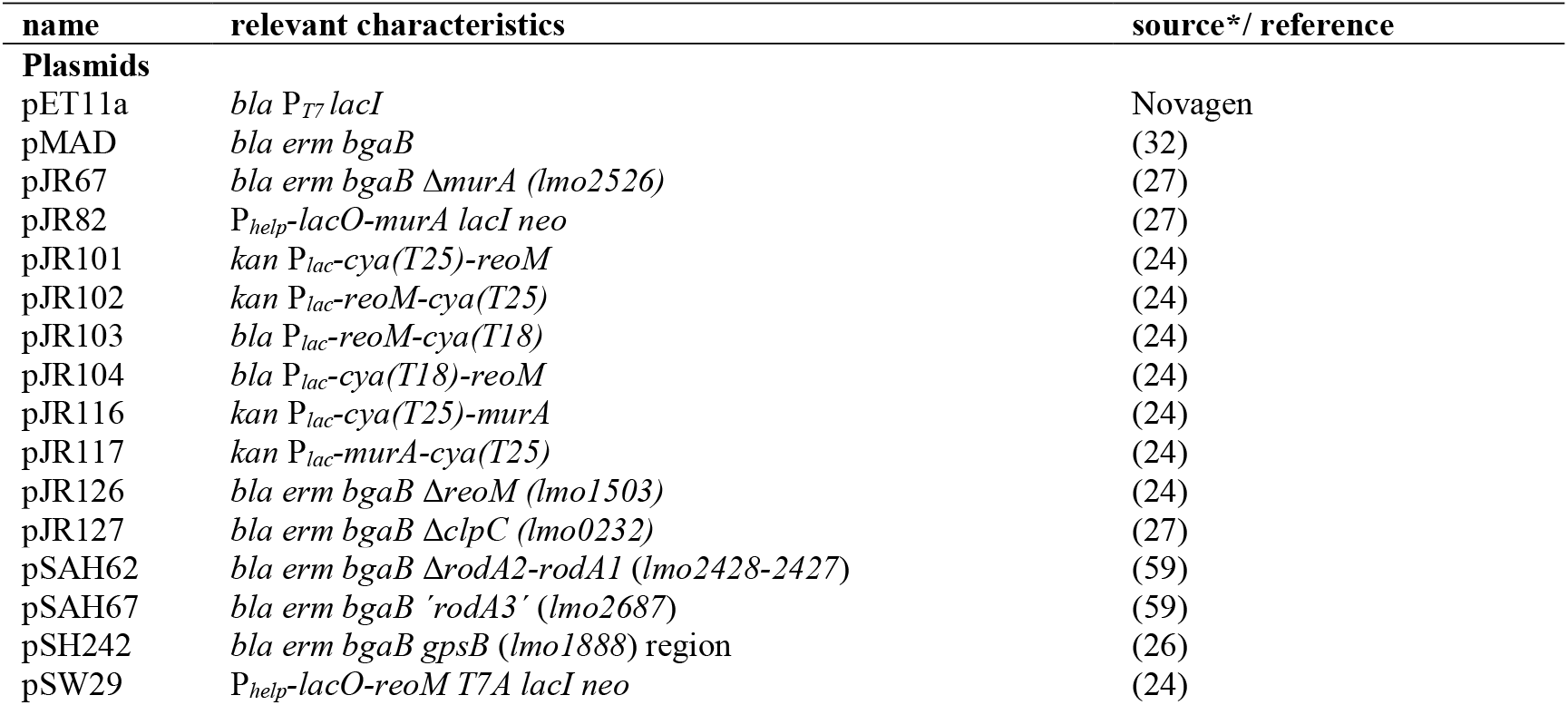
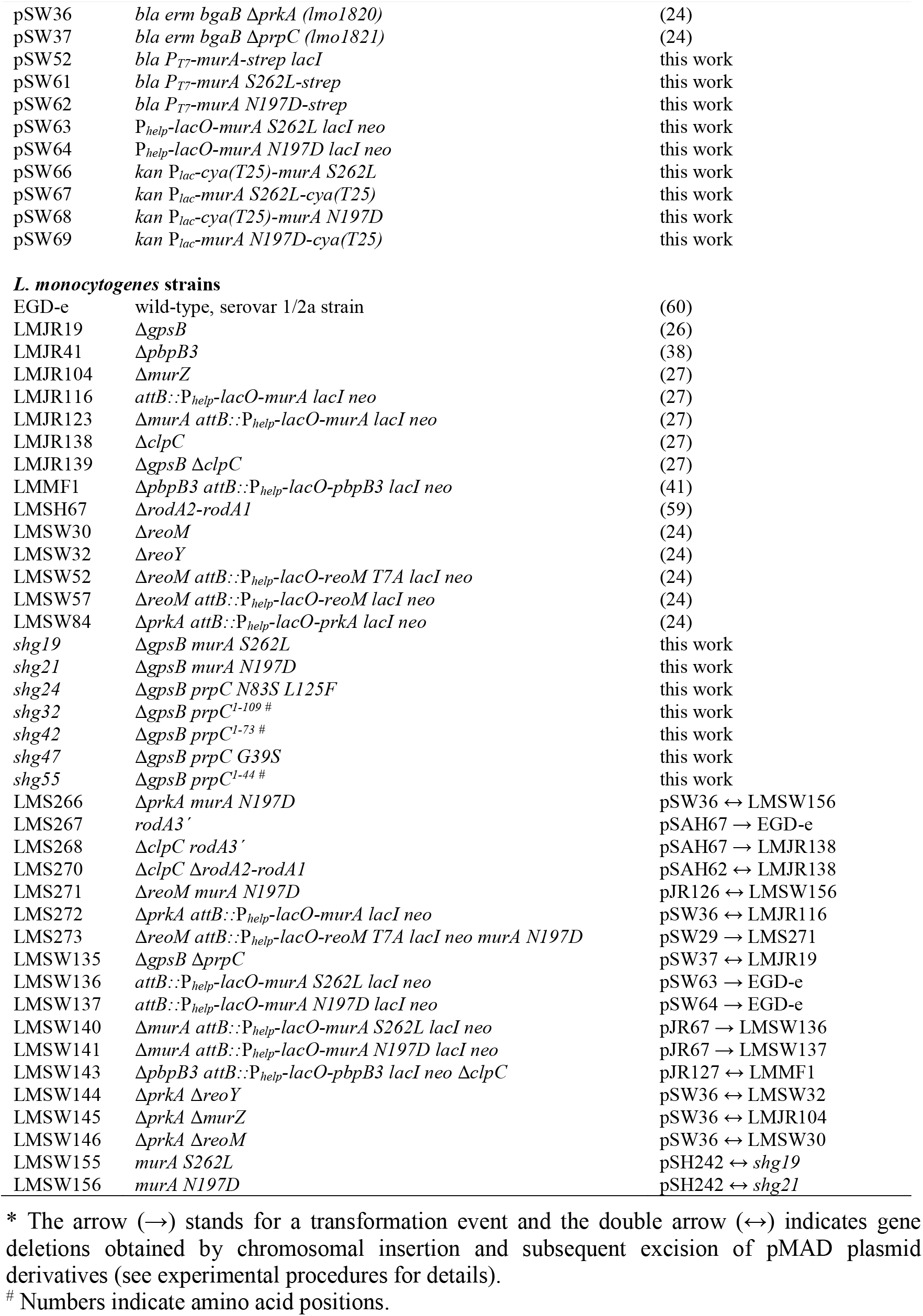
Plasmids and strains used in this study

### General methods, manipulation of DNA and oligonucleotide primers

Standard methods were used for transformation of *E. coli*, isolation of plasmid DNA and chromosomal DNA isolation (58). Transformation of *L. monocytogenes* was carried out by electroporation as described by others (61). The manufacturer’s instructions were followed for restriction and ligation of DNA. All primer sequences are listed in Tab 2.

**Table 2:**
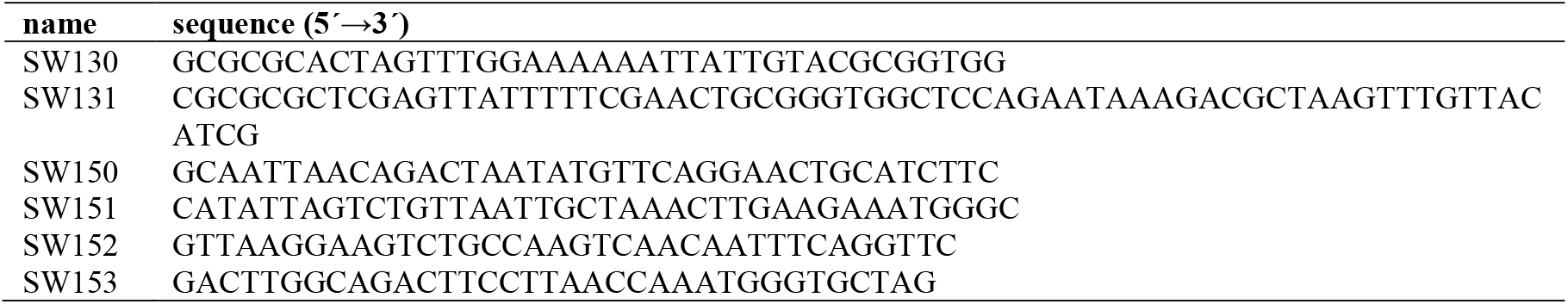
Oligonucleotides used in this study.

### Construction of plasmids and strains

Plasmid pSW52 was constructed for overexpression of Strep-tagged MurA in *E. coli*. To this end, the *murA* open reading frame was amplified with primers SW130/SW131 and cloned into pET11a using SpeI/XhoI. The S262L and N197D mutations were then introduced into pSW52 by quikchange mutagenesis using the primer pairs SW150/SW151 and SW152/SW153, respectively. The same primers were used to introduce both *murA* mutations into plasmid pJR82 by quickchange mutagenesis, yielding plasmids pSW63 and pSW64 for inducible expression of *murA S262L* and *murA N197D*, respectively, in *L. monocytogenes*. Likewise, these primers were used to introduce both mutations into the *murA* gene of the bacterial two hybrid vectors pJR116 and pJR117.

Derivatives of pIMK3 were introduced into *L. monocytogenes* strains by electroporation and clones were selected on BHI agar plates containing kanamycin. Plasmid insertion at the *attB* site of the tRNA^Arg^ locus was verified by PCR. Plasmid derivatives of pMAD were transformed into the respective *L. monocytogenes* recipient strains and genes were deleted as described elsewhere (32). All gene deletions and insertions were confirmed by PCR. Plasmid pSAH67, designed for insertional disruption of *rodA3*, was transformed into *L. monocytogenes* and transformants were selected on BHI agar plates containing erythromycin at 30°C. Plasmid insertion into the chromosomal *rodA3* gene was enforced by streaking the transformants to single colonies on BHI agar plates containing erythromycin at 40°C. Disruption of *rodA3* was confirmed by PCR.

### Bacterial two hybrid experiments

Plasmids carrying genes fused to T18- or the T25-fragments of the *Bordetella pertussis* adenylate cyclase were co-transformed into *E. coli* BTH101 (62) and transformants were selected on LB agar plates containing ampicillin (100 µg ml^-1^), kanamycin (50 µg ml^-1^), X-Gal (0.004%) and IPTG (0.1 mM). Agar plates were photographed after 48 h of incubation at 30°C.

### Genome sequencing

Chromosomal DNA for genome sequencing was isolated using the GenElute Bacterial Genomic DNA Kit (Sigma-Aldrich). Libraries were generated from 1 ng genomic DNA by using the Nextera XT DNA Library Prep Kit (Illumina). Sequencing was performed using a MiSeq Reagent Kit v3 cartridge (600-cycle kit) on a MiSeq benchtop sequencer in paired-end mode (2 × 300 bp). Reads were mapped to the *L. monocytogenes* EGD-e reference genome (NC_003210.1) (60) in Geneious (Biomatters Ltd.) to identify single nucleotide polymorphisms using the Geneious SNP finder tool. Genome sequences of *gpsB* suppressor strains were deposited at ENA under project accession number PRJEB47255.

### Isolation of cellular proteins and Western blotting

For protein isolation, cells from a 20 ml culture volume were harvested by centrifugation, washed with ZAP buffer (10 mM Tris.HCl pH7.5, 200 mM NaCl), resuspended in 1 ml ZAP buffer also containing 1 mM PMSF and disrupted by sonication. Cellular debris was removed by centrifugation and the supernatant was considered as total cellular protein extract. Aliquots of these protein samples were separated by SDS polyacrylamide gel electrophoresis and transferred onto positively charged polyvinylidene fluoride membranes employing a semi-dry transfer unit. DivIVA and MurA were detected by polyclonal rabbit antisera recognizing *B. subtilis* MurAA (63) and DivIVA (64) as the primary antibodies, respectively. An anti-rabbit immunoglobulin G conjugated to horseradish peroxidase was used as the secondary antibody and detected using the ECL chemiluminescence detection system (Thermo Scientific) in a chemiluminescence imager (Vilber Lourmat).

### Protein purification

Proteins were overproduced in *E. coli* BL21. Strains were cultivated in 1 l LB broth (containing 100 µg/ml ampicillin) at 37°C and 250 rpm. Protein expression was induced by addition of 1 mM IPTG at OD_600_ = 0.5. Cultivation was continued over night at 16°C and 200 rpm. Cells were harvested by centrifugation (11.325 x g, 10 min, 4°C) and the cell pellet was washed once in 20 ml ZAP buffer. Afterwards, the cell pellet was resuspended in 20 ml ZAP buffer containing 1 mM PMSF. Cells were disrupted using an EmulsiFlex C3 homogenizer (Avestin Europe GmbH) and cell debris was removed by centrifugation (4.581 x g, 10 min, 4°C) and the lysate was cleared in an additional passage through a filter (0.45 µm pore size). Strep-tagged proteins were purified using affinity chromatography and Strep-Tactin® Sepharose (IBA Lifesciences, Germany) according to the manufacturer’s instructions. Fractions containing purified proteins were pooled, aliquoted and stored at −20°C.

### MurA activity assay

An assay monitoring phosphate release from phosphoenol-pyruvate (PEP) was used for determination of MurA activity (65). For this purpose, 5 µg of purified MurA protein were mixed with 10 mM uridine 5′-diphospho-*N*-acetylglucosamine (UDP-Glc*N*Ac, Sigma-Aldrich) in a reaction buffer containing 100 mM Tris-HCl pH8.0 and 150 mM NaCl (final volume: 50 µl) and preincubated at 37°C for 15 min. The reaction was started by addition of 5 µl 10 mM PEP (Sigma-Aldrich) and stopped by addition of 800 µl staining solution after different time intervals. The staining solution was freshly prepared from 10 ml ammonium molybdate solution (4.2 g in 100 ml 4 M HCl), 30 ml malachite green solution (225 mg malachite green in 500 ml H_2_O) and 10 µl Triton X-100. Absorption was measured at λ=660 nm, corrected for background in the absence of UDP-Glc*N*Ac and used to calculate the amount of released phosphate using a standard curve generated with solutions with different phosphate concentrations.

### Lysis assays

Lysis assays were performed as described previously (66) with minor modifications. *L. monocytogenes* strains were grown in BHI broth at 37°C until an optical density of around OD_600_∼ 1.0. Cells were collected by centrifugation (6000 x g, 5 min, 4°C) and the cell pellet was resuspended in 50 mM Tris/HCl pH8.0 to an optical density of OD_600_ = 1.0. Lysozyme was added to a final concentration of 2.5 µg /ml (where indicated) and the cells were shaken at 37°C. Lysis was followed by measuring optical density every 15 min in a spectrophotometer.

### Macrophage infection assay

Experiments to measure intracellular growth of *L. monocytogenes* strains inside J774 mouse macrophages were essentially carried out as described earlier (67, 68).

### Microscopy

Cell membranes were stained through addition of 1 µl of nile red solution (100 µg ml^-1^ in DMSO) to 100 µl of exponentially growing bacteria. Images were taken with a Nikon Eclipse Ti microscope coupled to a Nikon DS-MBWc CCD camera and processed using the NIS elements AR software package (Nikon). Cell widths were determined by using the tools for distance measurements provided by NIS elements AR. Ultrathin section transmission electron microscopy was performed essentially as described earlier using a Tecnai 12 transmission electron microscope (Thermo Fisher/ FEI) operated at 120 kV (38).

## Acknowledgements

This work was funded by DFG grants HA 6830/1-2 and HA6830/4-1 (to S. H.). We would like to thank Birgitt Hahn for excellent technical assistance.

