## Supplementary figures S1-S6 for "MurA escape mutations uncouple peptidoglycan biosynthesis from PrkA signaling"

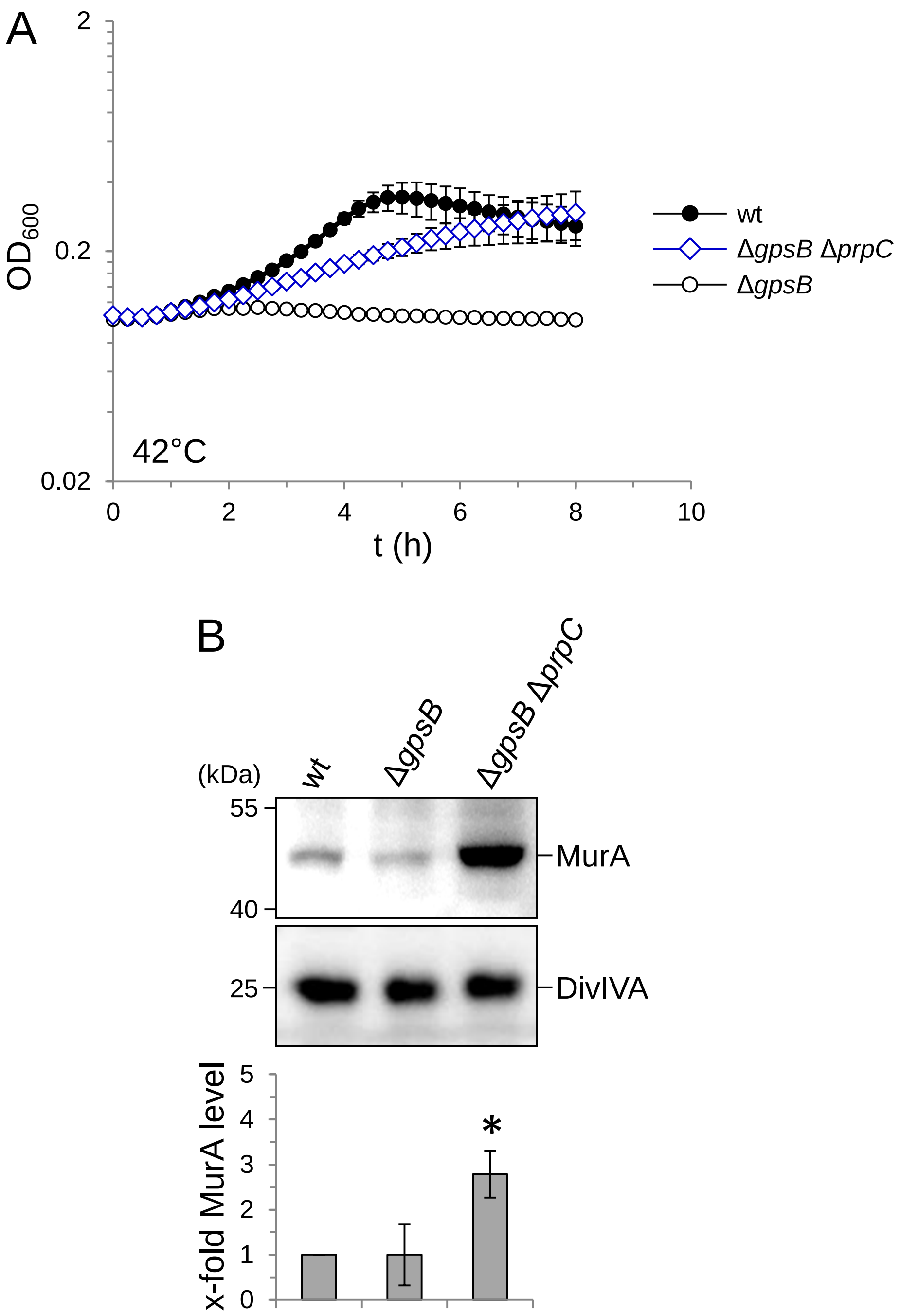


**Figure S1:** Suppression of the Δ*gpsB* phenotype by deletion of *prpC*.
(A) Growth of *L. monocytogenes* strains EGD-e (wt), LMJR19 (Δ*gpsB*) and LMSW135 (Δ*gpsB* Δ*prpC*) in BHI broth at 42°C. Average values and standard deviations were calculated from an experiment performed in triplicate.
(B) Western blot showing MurA and DivIVA levels (for control) in the same set of strains. MurA signals were quantified by densitometry and average values and standard deviations are shown (n=3). Asterisks indicated statistically significant differences (*t*-test, *P*<0.01).

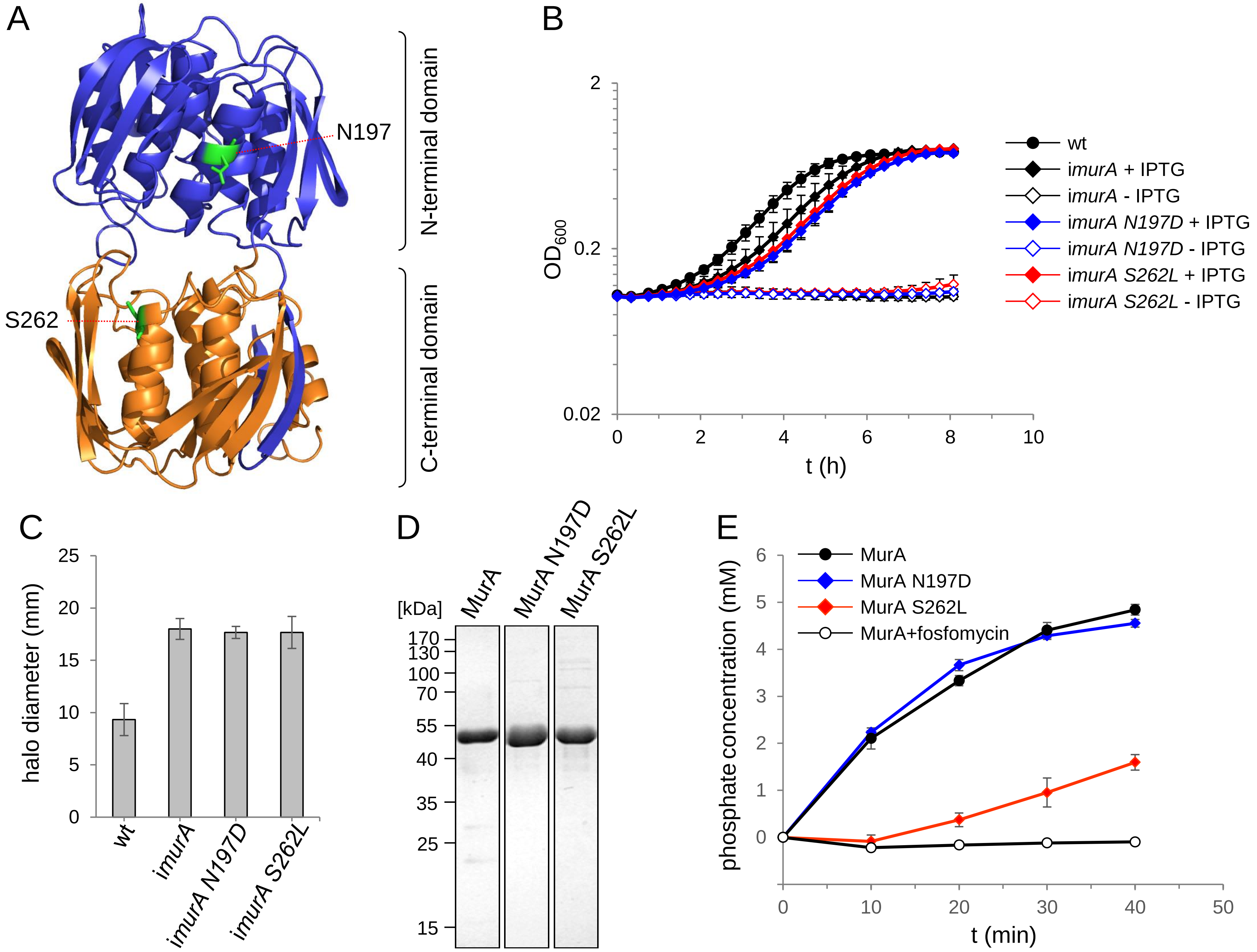


**Figure S2:** Effect of the N197D and S262L mutations on viability and MurA activity.

(A) Structural model of the *L. monocytogenes* MurA gene (PDB code: 3R38) (70) with the N197 and S262 residues indicated (green coloring).

(B) Effect of the *murA N197D* and *S262L* mutations on growth of *L. monocytogenes*. Strains EGD-e (wt), LMJR123 (i*murA*), LMSW140 (i*murA N197D*) and LMSW141 (i*murA S262L*) were grown in BHI broth ± 1 mM IPTG at 37°C. IPTG-dependent strains had to be pre-depleted during a growth passage in the absence of IPTG to develop fully visible IPTG-dependence. Average values and standard deviations from an experiment performed in triplicate are shown.

(C) Effect of the N197D and S262L mutations on fosfomycin susceptibility. The same strains as above were tested in a disc diffusion assay using filter discs soaked with fosfomycin on BHI agar plates not containing IPTG. The experiment was repeated three times and average values and standard deviations are shown.

(D) Purification of MurA-Strep and its N197D and S262L variants. Proteins were purified to near homogeneity and aliquots were separated using a standard SDS polyacryamid gel.

(E) Enzymatic activity of MurA-Strep, MurAN197D-Strep and MurAS262L-Strep. Average values and standard deviations calculated from three repetitions are shown.

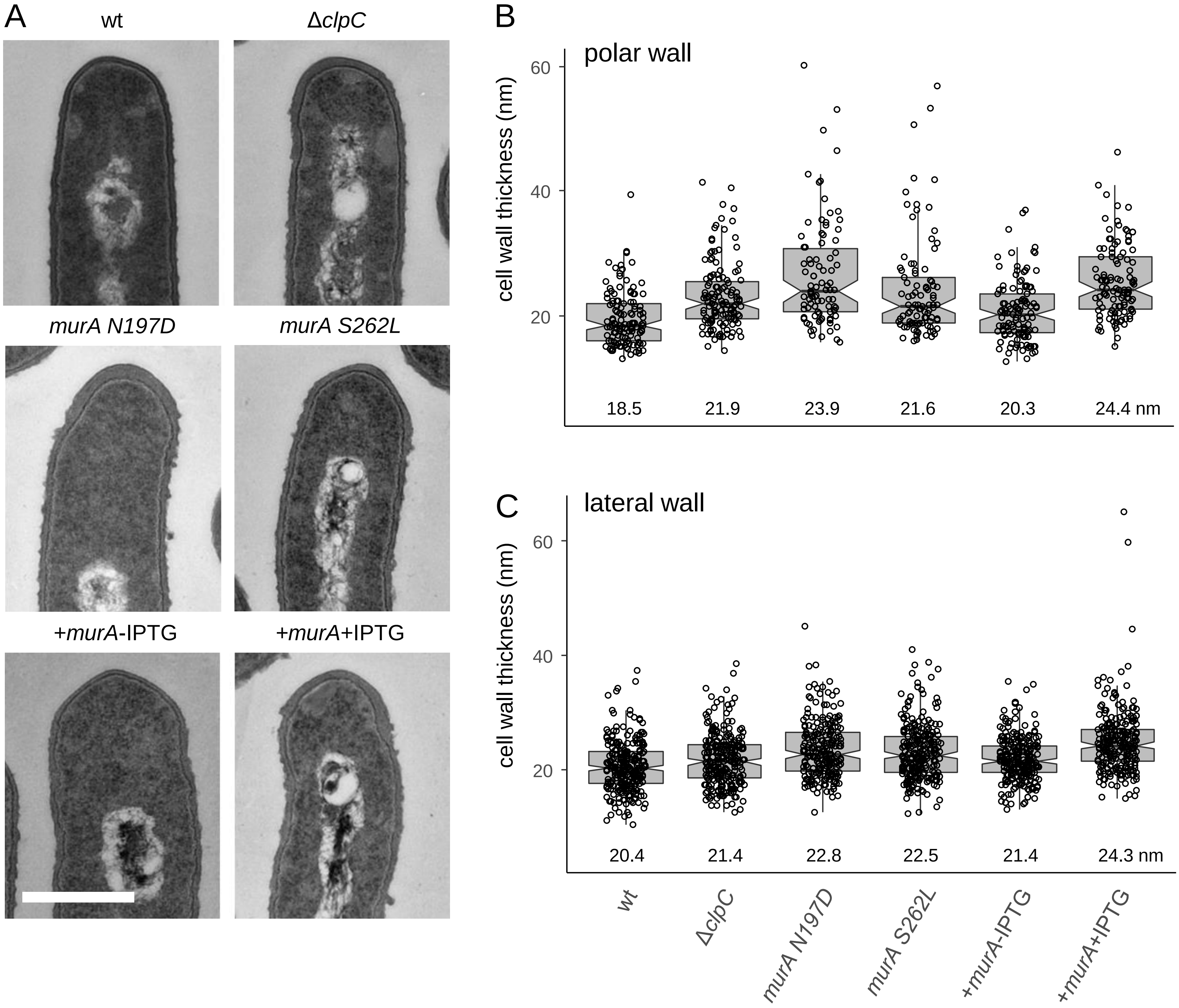


**Figure S3:** Thicker peptidoglycan in *murA* escape mutants

(A) Electron micrographs showing longitudinal sections of ultrathin-sectioned cells of *L. monocytogenes* strains EDG-e (wt), LMJR138 (Δ*clpC*), LMSW155 (*murA S262L*) and LMSW156 (*murA N197D*). Strains were grown in BHI broth at 42°C to early stationary phase ($OD_{600}$=1.5). Strain LMJR116 (+*murA*), which contains a second IPTG-inducible copy of *murA*, was included as control. Scale bar is 500 nm.

(B-C) Boxplots showing PG thickness at the cell poles (B) and the lateral wall (C). For determination of polar PG thickness, 16-21 longitudinally cut cells per strain were randomly selected and PG thickness was measured at three positions per pole, resulting in 96-166 measurements per strain. For determination of lateral PG thickness, 25 longitudinally cut cells per strain were selected and 10 measurements per cell were performed. Samples were blinded and mixed prior to the analysis. Median values are also shown.

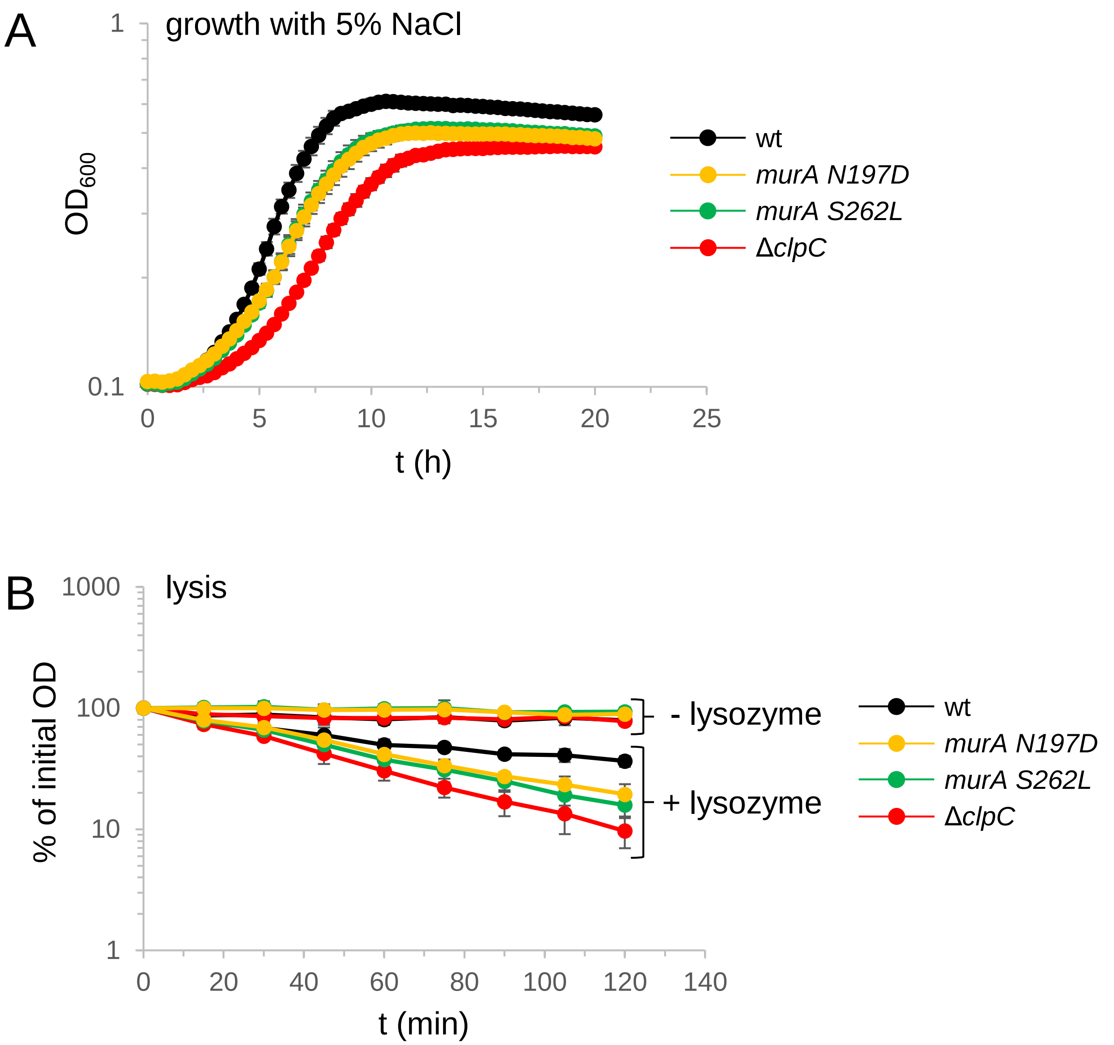


**Figure S4:** Sensitivity of *murA* escape mutants against salt and lysozyme.
(A) Growth of *L. monocytogenes* strains EDG-e (wt), LMJR138 (Δ*clpC*), LMSW155 (*murA S262L*) and LMSW156 (*murA N197D*) in BHI broth containing 5% (w/v) NaCl at 37°C. The experiment was performed in triplicate and average values and standard deviations are shown.
(B) Lysis of the same set of strains in the presence of lysozyme. The experiment was performed three times, average values and standard deviations are shown.

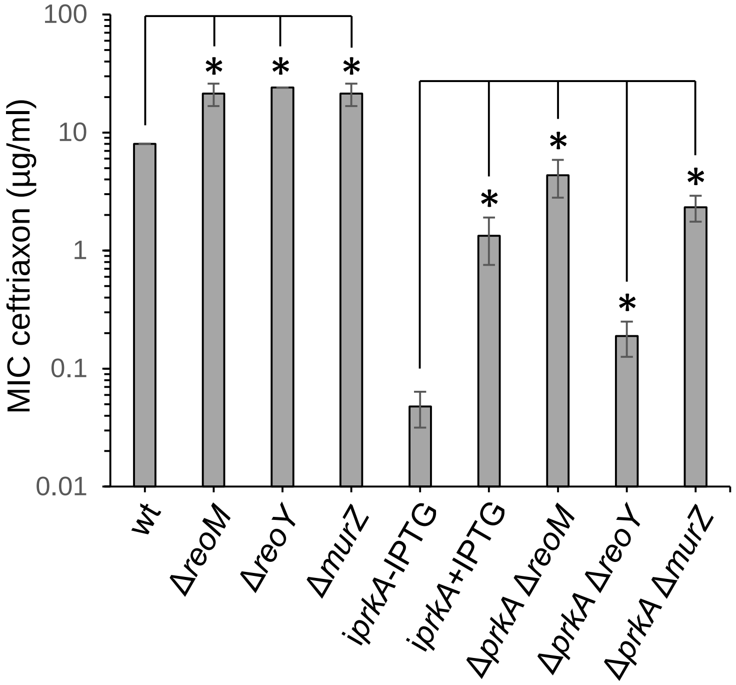


**Figure S5:** Suppression of the *prkA* phenotype by *reoM*, *reoY* and *murZ* deletions
Minimal inhibitory ceftriaxone concentrations of *L. monocytogenes* strains EGD-e (wt), LMSW30 (Δ*reoM*), LMSW32 (Δ*reoY*), LMJR104 (Δ*murZ*), LMSW84 (i*prkA*), LMSW146 (Δ*prkA* Δ*reoM*), LMSW144 (Δ*prkA* Δ*reoY*) and LMSW145 (Δ*prkA* Δ*murZ*) are shown. Values represent average values from three repetitions. Asterisks mark statistical significance ($P$<0.05, *t*-test).

| | PASTA-eSTK | PrpC | ReoM | ReoY | MurA1 | MurA2 | RodA3 | PBP B3 |
|---|---|---|---|---|---|---|---|---|
| ***Lmo*** | PrkA<br>*lmo1820* | PrpC<br>*lmo1821* | ReoM<br>*lmo1503* | ReoY<br>*lmo1921* | MurA<br>*lmo2526* | MurZ<br>*lmo2552* | RodA3<br>*lmo2687* | PbpB3<br>*lmo0441* |
| ***Bsu*** | PrkC<br>BSU_15770<br>(0.0) | PrpC<br>BSU_15760<br>(7e-82) | ReoM<br>BSU_27400<br>(1e-39) | ReoY<br>BSU_22580<br>(4e-61) | MurAA<br>BSU_36760<br>(0.0) | MurAB<br>BSU_37100<br>(0.0) | RodA<br>BSU_38120<br>(1e-40) | PbpC<br>BSU_04140<br>(e-140) |
| ***Efa*** | IreK<br>EF3120<br>(2e-176) | IreP<br>EF3121<br>(3e-77) | IreB<br>EF1202<br>(1e-33) | ReoY<br>EF1554<br>(3e-39) | MurAA<br>EF2605<br>(0.0) | MurAB<br>EF1169<br>(0.0) | RodA<br>EF2502<br>(5e-57) | Pbp4<br>EF2476<br>(0.0) |
| ***Sau*** | Stk1<br>SA1063<br>(e-143) | Stp1<br>SA1062<br>(3e-61) | ReoM<br>SA1445<br>(2e-26) | ReoY<br>SA1295<br>(3e-08) | MurA<br>SA1902<br>(0.0) | MurZ<br>SA1926<br>(e-166) | RodA<br>SA1888<br>(9e-48) | MecA<br>SA0038<br>(e-121) |
| ***Spn*** | StkP<br>SPR1577<br>(4e-145) | PhpP<br>SPR1578<br>(6e-55) | ReoM<br>SPR0175<br>(2e-29) | - | MurA<br>SPR1781<br>(0.0) | MurZ<br>SPR0989<br>(3e-155) | RodA<br>SPR0712<br>(9e-51) | - |

E value

- x<e-100
- e-100<x<e-20
- x>e-20

**Figure S6:** Conservation of the PrkA signaling cascade in selected Gram-positive bacteria.
Components of the PrkA signaling route in *L. monocytogenes* EGD-e (*Lmo*) and their homologues in *B. subtilis* 168 (*Bsu*), *E. faecalis* V583 (*Efa*), *S. aureus* N315 (*Sau*) and *S. pneumoniae* R6 (*Spn*). Locus numbers are given below the protein names and protein sequence homologies are shown as e-values (in brackets).
